# A dynamic spatiotemporal normalization model for continuous vision

**DOI:** 10.1101/2025.03.06.641906

**Authors:** Angus F. Chapman, Rachel N. Denison

**Author notes:** Corresponding Author: Angus Chapman. Author contributions: Conceptualization: A.F.C., R.N.D. Methodology: A.F.C., R.N.D. Software: A.F.C., R.N.D. Formal Analysis: A.F.C. Visualization: A.F.C. Supervision: R.N.D. Writing – Original Draft: A.F.C. Writing – Review & Editing; A.F.C., R.N.D.

## Abstract

How does the visual system process dynamic inputs? Perception and neural activity are shaped by the spatial and temporal context of sensory input, which has been modeled by divisive normalization over space or time. However, theoretical work has largely treated normalization separately within these dimensions and has not explained how future stimuli can suppress past ones. Here we introduce a dynamic spatiotemporal normalization model (D-STAN) with a unified spatiotemporal receptive field structure that implements normalization across both space and time and ask whether this model captures the bidirectional effects of temporal context on neural responses and behavior. D-STAN implements temporal normalization through excitatory and suppressive drives that depend on the recent history of stimulus input, controlled by separate temporal windows. We found that biphasic temporal receptive fields emerged from this normalization computation, consistent with empirical observations. The model also reproduced several neural response properties, including surround suppression, nonlinear response dynamics, subadditivity, response adaptation, and backwards masking. Further, spatiotemporal normalization captured bidirectional temporal suppression that depended on stimulus contrast, consistent with human behavior. Thus, D-STAN captured a wide range of neural and behavioral effects, demonstrating that a unified spatiotemporal normalization computation could underlie dynamic stimulus processing and perception.

## Introduction

Although the world around us is highly dynamic, most theories and models of visual processing consider only a static snapshot of this constantly changing stream of visual stimulation. Less understood is the dynamic neural activity that supports visual perception, which exhibits several non-linear response properties that depend on time-varying stimulus input. Neurons in a range of cortical areas show sensitivity to time-varying stimuli (Mauk & Buonomano, 2004), such as motion-sensitive neurons in V1 and MT, which can be characterized by their *spatiotemporal* receptive fields. More broadly, many neural responses and perception depend not just on current inputs but also on recent stimulus history (Fritsche et al., 2020; Gao et al., 2020; Hasson et al., 2008; Murray et al., 2014; Wolff et al., 2022), and can be modulated by future context as well (Breitmeyer & Öğmen, 2006; Yeshurun et al., 2015). These findings demonstrate that sensory systems are sensitive to temporal structure, which raises questions about what mechanisms exist to process dynamic inputs (Cavanagh et al., 2020; Golesorkhi et al., 2021; Soltani et al., 2021).

A powerful theoretical framework for visual processing is based on the principle of normalization, which has been proposed as a canonical neural computation (Carandini & Heeger, 2012). Normalization is the idea that neurons can have suppressive effects on one another as a function of their tuning preferences, acting to “normalize” overall activity levels within a neural population (Heeger, 1992). Most normalization models are static, with the normalization computation operating across cortical space, which we refer to as spatial normalization. Temporal normalization, in contrast, is the idea that neural activity undergoes normalization across time (Heeger, 1992). Current dynamic normalization models implement temporal normalization by using a temporally filtered and delayed version of the excitatory input drive to compute the normalization signal (Groen et al., 2022; Zhou et al., 2019). Other models combine linear stimulus evoked responses with non-linear compressive—rather than divisive— computations to model how neurons respond to dynamic input (Kim et al., 2024; Kupers et al., 2024). All these models better predict neural response dynamics than linear-only models and capture several non-linear phenomena (Zhou et al., 2019). However, they also have important limitations. First, the entire excitatory time course must be known in advance, limiting their biological feasibility. Second, normalizing by a delayed copy of the excitatory drive means that only past stimuli can suppress future stimuli, not vice versa. Third, normalization is computed for each neural unit (neuron, voxel, electrode, etc.) in isolation, so to date these models have not considered how temporal normalization may operate across a broader suppressive pool, which has been critical for the success of spatial normalization models.

Here we introduce the dynamic spatiotemporal attention and normalization model (D-STAN), a model of visual processing in which spatial and temporal normalization are integrated within a unified receptive field-based spatiotemporal normalization computation. Model neurons are situated in a neural network architecture, performing real-time processing of continuous visual inputs. We analyzed the response properties of this model, focusing on its time-varying behavior and modulation by temporal context. By unifying normalization across space and time, we find that D-STAN captures key non-linear temporal response properties of neurons and predicts human behavior. Critically, unlike previous dynamic normalization models, it can produce bidirectional temporal suppression between stimuli in a sequence, allowing for empirically demonstrated effects of temporal context that operate both forward and backward in time. The current work builds on the normalization model of dynamic attention developed by Denison et al. (2021), which generalized the normalization model of attention (Reynolds & Heeger, 2009) to the time domain. D-STAN advances previous work by implementing excitatory and suppressive drives that depend on recent stimulus history, imbuing the model neurons with receptive fields and normalization computations that are inherently spatiotemporal.

## Results

### The dynamic spatiotemporal attention and normalization model (D-STAN)

We introduce a model of dynamic visual perception and attention building on (Denison et al., 2021) that produces neural responses and behavioral outputs. The model simulates sensory responses to stimulus input through neurons that are tuned to specific spatial locations and feature values and, critically, integrate inputs over the recent past, giving them spatiotemporal receptive fields (Figure 1A). As in previous models, spatial and feature dimensions are treated equivalently (Reynolds & Heeger, 2009), with normalization across these dimensions determined by the tuning properties of model neurons. Sensory responses can also be modulated by time-varying attention, though we do not explore attentional modulation here. The sensory responses are read out by a decision layer, which accumulates evidence toward a particular behavioral output. Importantly, we include spatiotemporal normalization at each stage of processing, with excitatory and suppressive drives that contribute to the final neural response (Figure 1B). Responses are continuously updated at each time point with differential equations, allowing us to examine how model parameters affect the dynamics of neural responses as well as the final behavioral output. Thus, the model generates neural responses continuously at each time step (Figure 1C), at a level of abstraction that allows us to focus on how excitatory and suppressive components affect population activity separate from single unit physiology or neural circuit dynamics.

**Figure 1.**
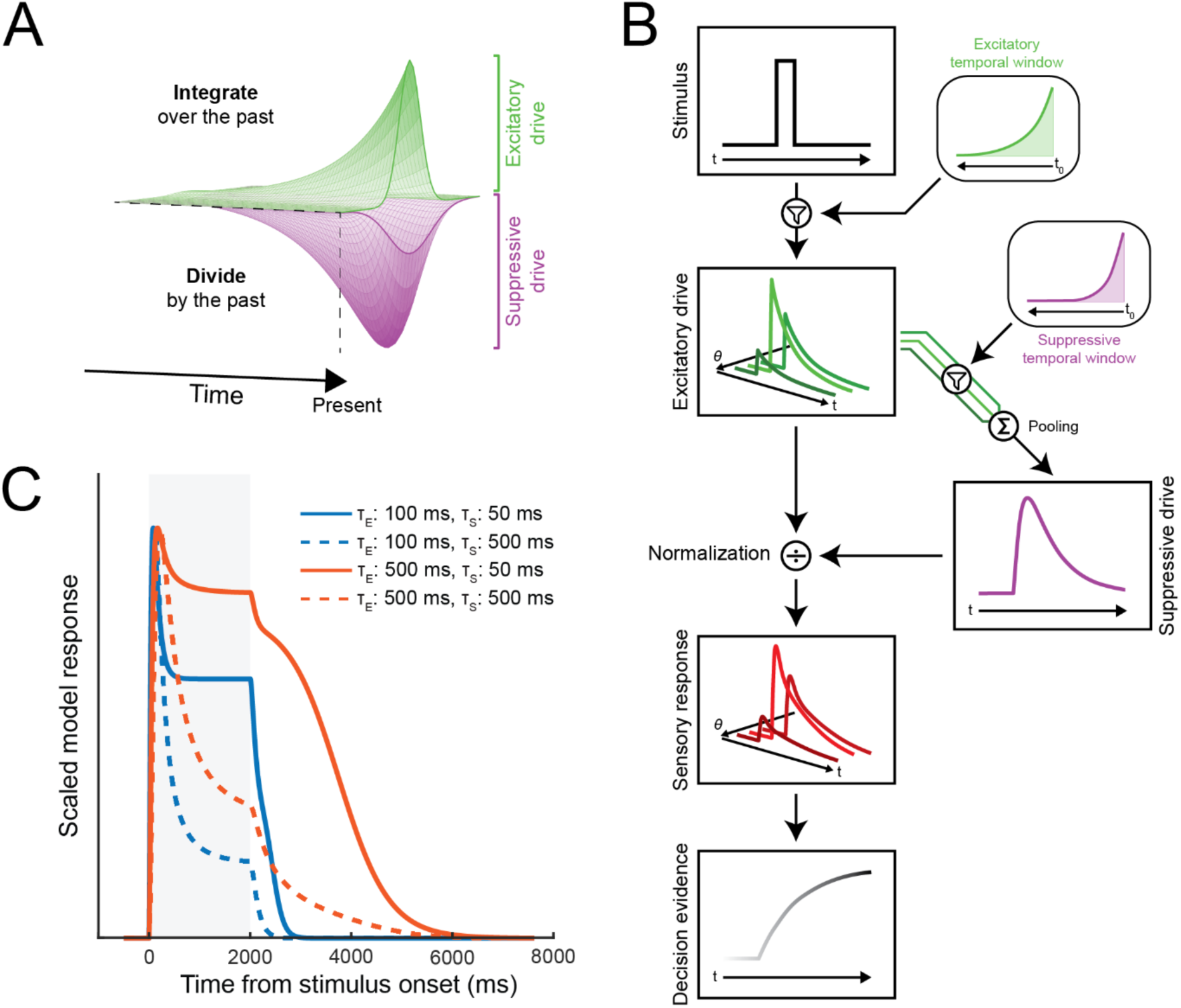
Architecture of the Dynamic Spatiotemporal Attention and Normalization model (D-STAN). A) Spatiotemporal receptive field structure. Model sensory neurons integrate over the recent history of stimulus input, while contributing to a suppressive pool that also extends into the past. Model neurons can be tuned to different spatiotemporal properties of the stimulus input as well as to other stimulus features. B) Schematic of computations in D-STAN. The model receives time-varying stimulus input (here, an oriented stimulus with a specific contrast), which is continuously filtered by the excitatory temporal window. The stimulus input produces excitatory drives in model neurons tuned to different orientations (θ). The excitatory drives of each neuron are continuously filtered and pooled to calculate the suppressive drive, which in turn normalizes the excitatory drives to compute sensory responses. Sensory layer activity is fed into the decision layer, which accumulates evidence about the stimulus orientation. C) Example sensory layer responses to a stimulus presentation (shaded grey region) as a function of different excitatory and suppressive temporal window parameters.

The core of the model is the spatiotemporal normalization computation, which divides the spatiotemporal excitatory drive by the spatiotemporal suppressive drive. To investigate the effects of temporal normalization, we implemented temporal receptive fields in model sensory neurons by allowing the excitatory drive to depend on stimulus input at previous points in time. Specifically, at each time point, the stimulus time course was weighted by an exponential decay function, such that input at more distant points in the past had less effect on the neuron’s response. The time constant (τ_E_) determined the relative weight given to stimuli at each point in time. When τ_E_ is zero, the model neuron only receives excitatory drive when stimulus input is present, but increasing τ_E_ results in responses that are driven even after the stimulus ends. Additionally, the suppressive drive of the neural population was pooled across neural units, as in previous work (Denison et al., 2021; Reynolds & Heeger, 2009), but also across time. The suppressive drive was calculated by weighting the excitatory drive across previous time points by an exponential decay function with a separate time constant (τ_S_). The suppressive drive is thus dependent on both time constants with the effect that suppression always acts later and across longer time intervals than excitation (even when τ_S_ < τ_E_). We performed a series of simulations varying aspects of the stimulus input to sensory layers to examine whether the model could reproduce several different non-linear response properties and to determine the effect of the excitatory and suppressive time constants on these response dynamics.

### Stimulus history differentially affects model drives and responses

The dependence of a neuron’s current response on recent stimulus inputs defines its temporal receptive field. We used reverse correlation to determine how excitatory and suppressive temporal windows interact to shape the functional temporal receptive fields of model neurons. We first examined how the model responses depended on stimulus history for a specific set of excitatory and suppressive temporal windows (τ_E_ = 400 ms, τ_S_ = 100 ms). Random stimulus input was fed into the model sensory layer, driving variable activity across simulations. We calculated how the presence of stimuli up to 1200 ms in the past affected model neuron responses at the current moment by correlating the random stimulus vectors with the observed excitatory drive, suppressive drive, and sensory layer response. As expected, the excitatory drive depended most strongly on input close to the current time, with weights back in time following the exponential excitatory temporal window (Figure 2A). The suppressive drive depended on times in the recent past, and could be approximated by a convolution of the excitatory and suppressive temporal windows (Figure 2B), consistent with how suppressive drives are a temporally-weighted sum of all model neurons’ excitatory drives, meaning both τ_E_ and τ_S_ affect the shape of the suppressive drive.

**Figure 2.**
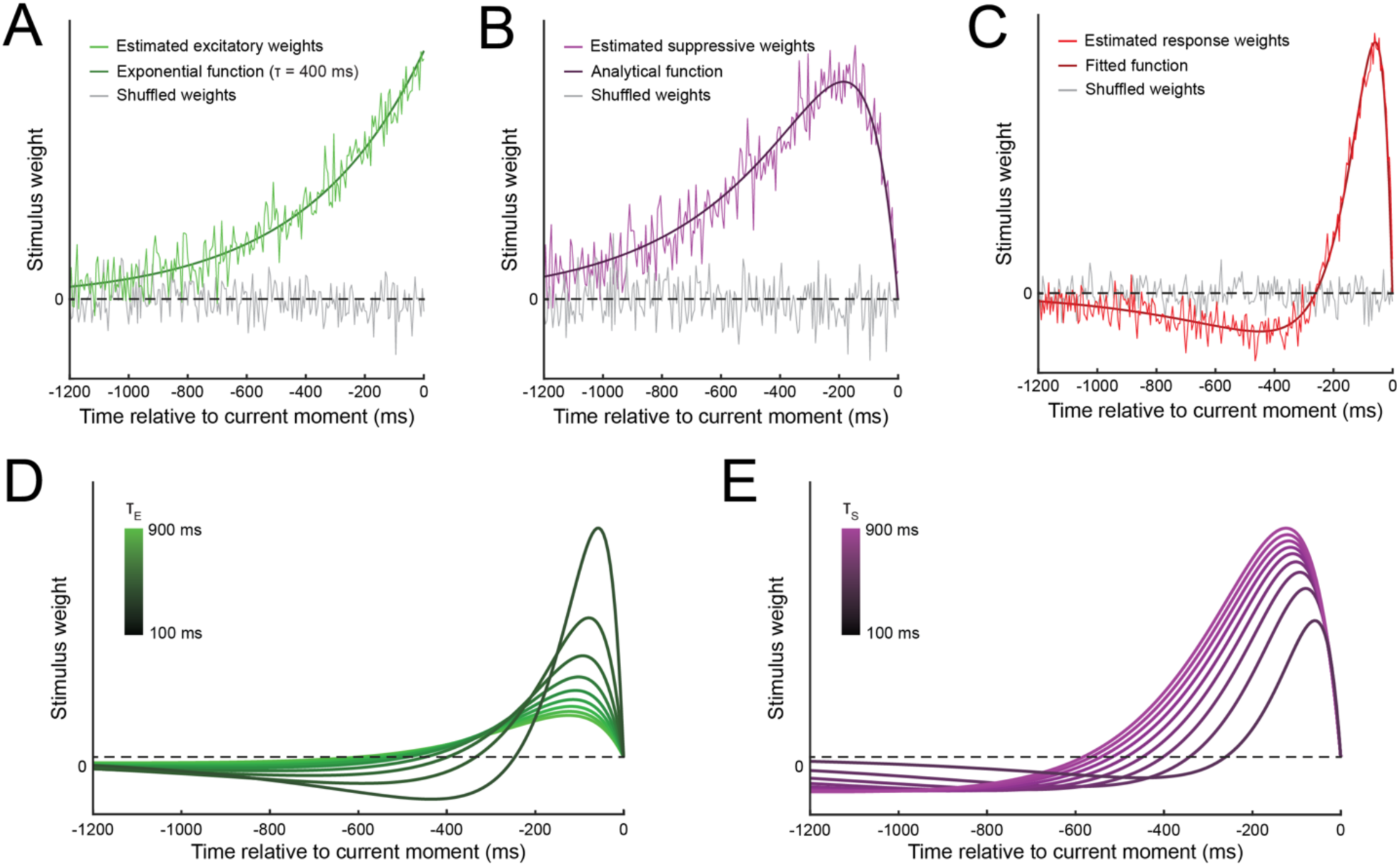
Reverse correlation analysis of model drives and sensory response reveals biphasic temporal receptive field. Stimulus weights were estimated for A) excitatory drives, B) suppressive drives, and C) sensory layer responses. Examples shown reflect simulations with τ_E_ = 400 ms and τ_S_ = 100 ms. Each set of estimated weights were fitted using different functional forms, (see main text). We also estimated the sensory layer weight functions (i.e., temporal receptive fields) across a range of parameters for D) excitatory temporal windows, and E) suppressive temporal windows. While varying one parameter, the other was fixed at an intermediate level (400 ms) for visualization purposes; the pattern of results did not depend on the specific fixed value.

Reverse correlation revealed that this implementation of normalization generates a biphasic temporal receptive field (Figure 2C), where stimulus input close to the current time drives model neuron responses and input further in the past reduces responses. Notably, the biphasic stimulus weighting function was not built directly into the model, but emerged through the interaction between the excitatory and suppressive temporal windows. A similar biphasic function in time has been found in empirical recordings from visual neurons (Cai et al., 1997; Mante et al., 2008; Perge et al., 2005), and is similar to those used in models of neural temporal dynamics (Kim et al., 2024; Kupers et al., 2024; Zhou et al., 2019). The estimated response weights were well fitted by a difference of Gamma functions, following previous work (Mante et al., 2008; Zhou et al., 2019).

We performed additional simulations to explore the effect of the excitatory and suppressive time constants on the shape of the temporal receptive fields. Increasing the excitatory time constant, while holding the suppressive time constant fixed (τ_S_ = 400 ms), resulted in a scaling and extension of the temporal receptive field to times further in the past (Figure 2D), as stimuli at these times fell into the longer excitatory temporal windows—a “flattening out” of the response profile. Increasing the suppressive time constant, with the excitatory time constant fixed (τ_E_ = 400 ms), also resulted in an extension of the temporal receptive field to past times (Figure 2E), although this was a consequence of the suppression being distributed over a longer time span, resulting in less suppression for more recent times. Increasing the suppressive time constant also resulted in longer periods of suppression that extended further into the past, as would be expected given the longer suppressive windows. Thus, the excitatory and suppressive temporal windows generated temporal receptive fields that exhibit a variety of response profiles depending on each time constant.

### Reproducing signatures of spatial normalization

Before turning to the time-dependent aspects of the model, we sought to ensure that D-STAN could capture effects associated with static spatial normalization as expected. One finding commonly attributed to normalization is surround suppression: the suppression of a neuron’s response when a competing stimulus is placed in the region surrounding its excitatory receptive field (Figure 3A) (Cavanaugh et al., 2002a, 2002b). The explanation given by spatial normalization is that while the neuron may not be responsive to stimuli placed in the surround region alone, other neurons that are tuned to stimuli in those spatial locations contribute to the normalization pool, such that activity driven by surrounding stimuli suppresses neurons tuned to center stimuli. We reproduced this effect in D-STAN by varying the contrast of the center and surround stimuli independently, following (Carandini, 2004; Carandini & Heeger, 2012). The model responses exhibited sigmoidal contrast response functions, as is typical of normalization models (Carandini et al., 1997; Heeger, 1992). While the model response to the center stimulus increased monotonically as a function of contrast, its response was simultaneously suppressed by a stimulus in the surround, with stronger suppression for higher surround contrasts (Figure 3B). Thus, D-STAN captures a key signature of spatial normalization.

**Figure 3.**
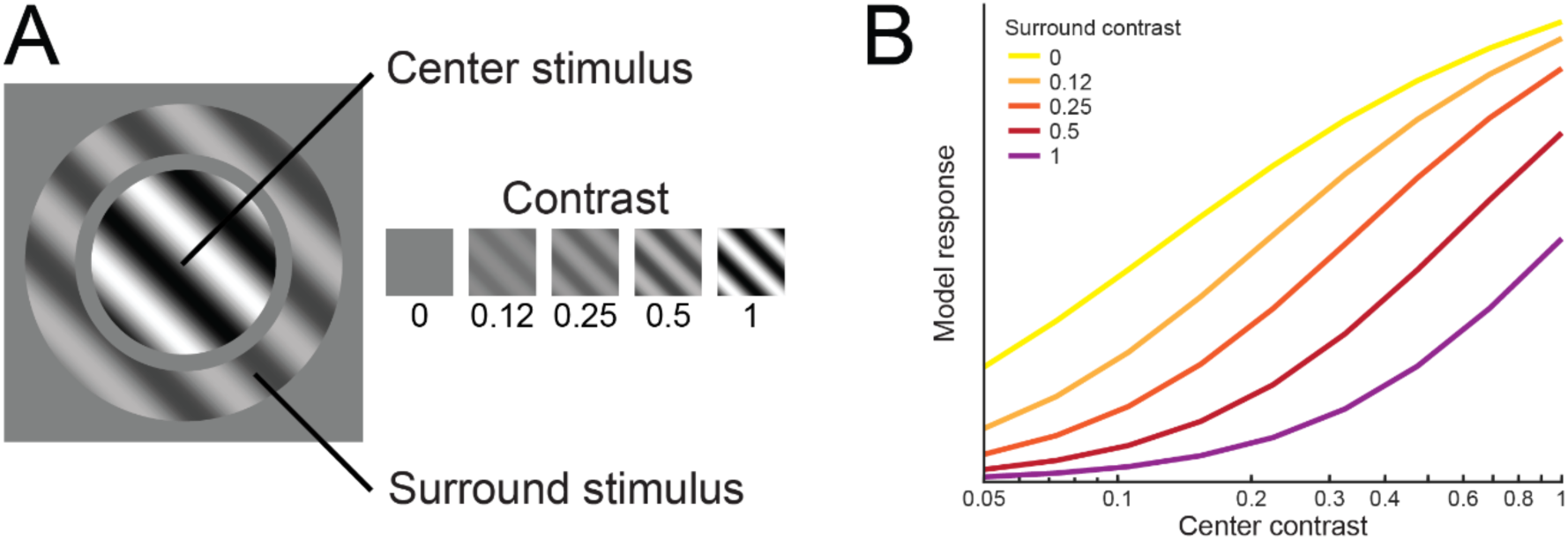
Model sensory neurons exhibit contrast-dependent surround suppression, a signature of spatial normalization. A) We measured the response of neurons tuned to the orientation and spatial location of the center stimulus while manipulating the contrast of both the center and surround stimuli (in this example center contrast = 1, while surround contrast = 0.5). The center and surround were presented simultaneously to model neurons for 100 ms, and model temporal window parameters were fixed across simulations (τ_E_ = 400 ms, τ_S_ = 100 ms). B) Responses showed the expected contrast response function as the center stimulus contrast increased, as well as increasing suppression from the surround as its contrast increased.

### Reproducing known temporal properties of neuronal responses

We next asked whether D-STAN could reproduce several known non-linear temporal response properties of neurons: 1) transient-sustained dynamics, 2) subadditivity, and 3) response adaptation. For each property we asked whether excitatory temporal windows, suppressive temporal windows, or both were necessary to reproduce the property and how the non-linear temporal effects depended on the excitatory and suppressive time constants in the model.

#### Transient-sustained dynamics

Neurons typically show an initial transient response followed by sustained activity to a prolonged stimulus (Lisberger & Movshon, 1999; Motter, 2006). We found that the suppressive temporal window was necessary to reproduce these dynamics. When varying the excitatory time constant with no suppressive temporal window (τ_S_ = 0), responses increased gradually towards a stable activity level following stimulus onset and decreased only after stimulus offset, consistent with linear predictions (Zhou et al., 2019). Higher τ_E_ resulted in slower rise times (Figure 4A) and consequently longer times to peak (Figure 4B), as well as more prolonged responses after stimulus offset (Figure 4C). The model exhibited slower response dynamics with increasing τ_E_, because longer integration windows provided excitatory drive from stimulus input further in the past. In contrast, when varying the suppressive time constant alone (τ_E_ = 0), model responses showed a more typical transient response, with an initial peak shortly after stimulus onset that decreased to a stable level before stimulus offset (Figure 4D). Higher τ_S_ resulted in longer times to peak (Figure 4E) and a lower stable activity level relative to peaks (Figure 4F), because suppression integrated more slowly but reached greater levels overall. Varying both time constants generated model responses with a diverse range of temporal profiles (Figure 1C). Thus, suppression alone, or a combination of excitatory and suppressive temporal windows, can capture typical neural dynamics to prolonged stimulus presentations.

**Figure 4.**
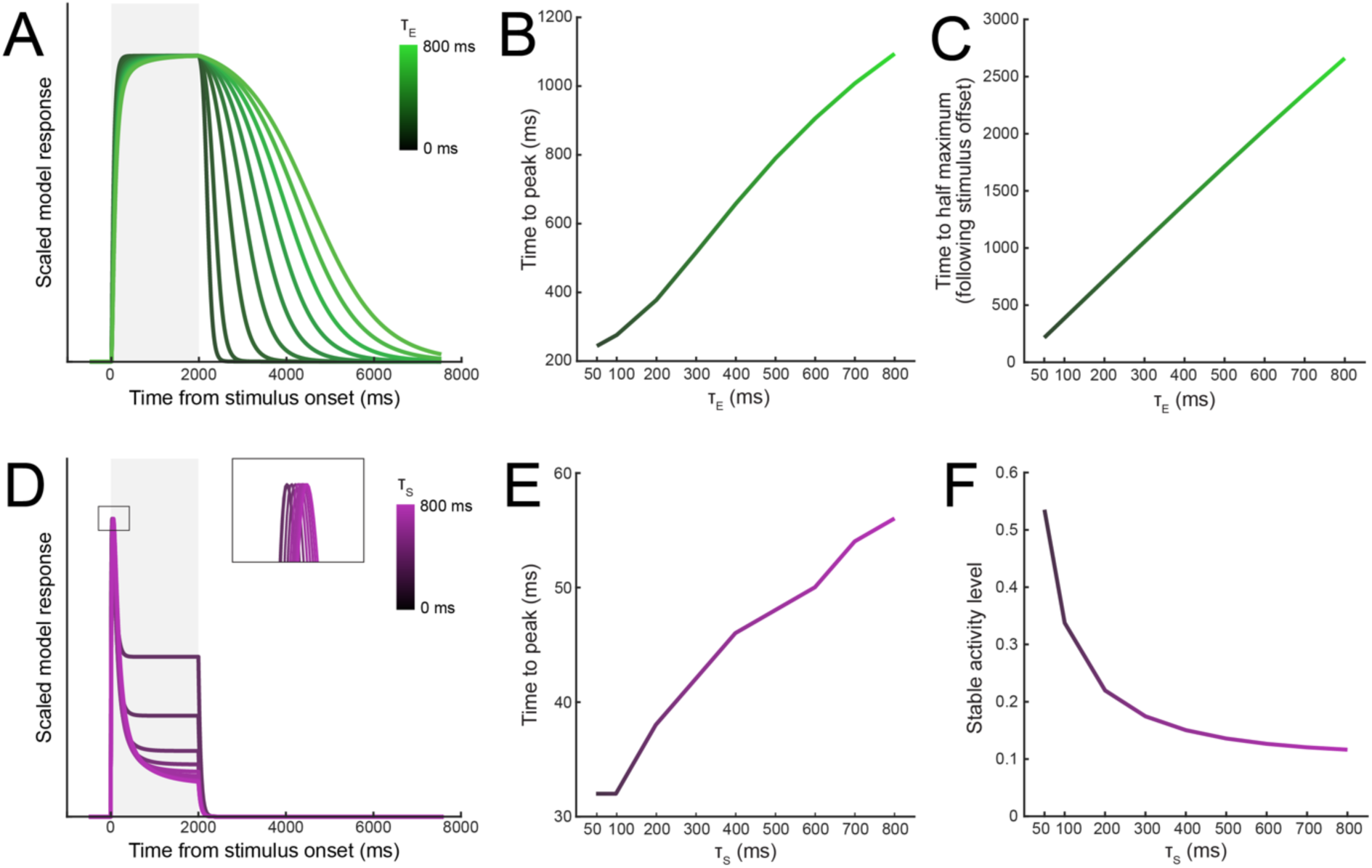
Excitatory and suppressive time constants differentially contribute to transient-sustained response dynamics. A) Effect of the excitatory time constant, ranging from 0-800 ms in 100 ms steps. Shaded region shows the stimulus presentation period. B) Time to peak for sensory responses as a function of τ_E_. C) Time for sensory responses to reduce to 50% of maximum as a function of τ_E_. D) Effect of the suppressive time constant on model sensory responses. The insert shows the early peak responses. E) Time to peak for the sensory responses as a function of τ_S_. F) Stable activity level of sensory responses, relative to the peaks, reached by the end of the stimulus presentation period as a function of τ_S_.

#### Subadditivity

Another hallmark of non-linear response dynamics is that neural responses display temporal subadditivity: doubling the duration of the stimulus leads to a less than doubling of the response amplitude (Groen et al., 2022; Zhou et al., 2018). Increasing the stimulus duration in D-STAN resulted in more prolonged model responses (Figure 5A) that were strongly subbadditive, as quantified by the area under each curve (Figure 5B). We found that increasing either τ_E_ or τ_S_ was sufficient to produce subadditive model responses (Figure 5C-D); when both time constants were zero, model responses fell just below the linear prediction, showing only slight subadditivity due to the time constant inherent to the model’s recursive computation (τ_R_, Equation 6). For values of τ_E_ and τ_S_ above zero, each doubling of the stimulus duration increased responses by only ∼1.3-1.6×. Combining different parameter values produced similar levels of subadditivity (Figure 5E), where lower values of each time constant resulted in relatively more subadditivity at short stimulus durations, with higher values resulting in more subadditivity at longer stimulus durations.

**Figure 5.**
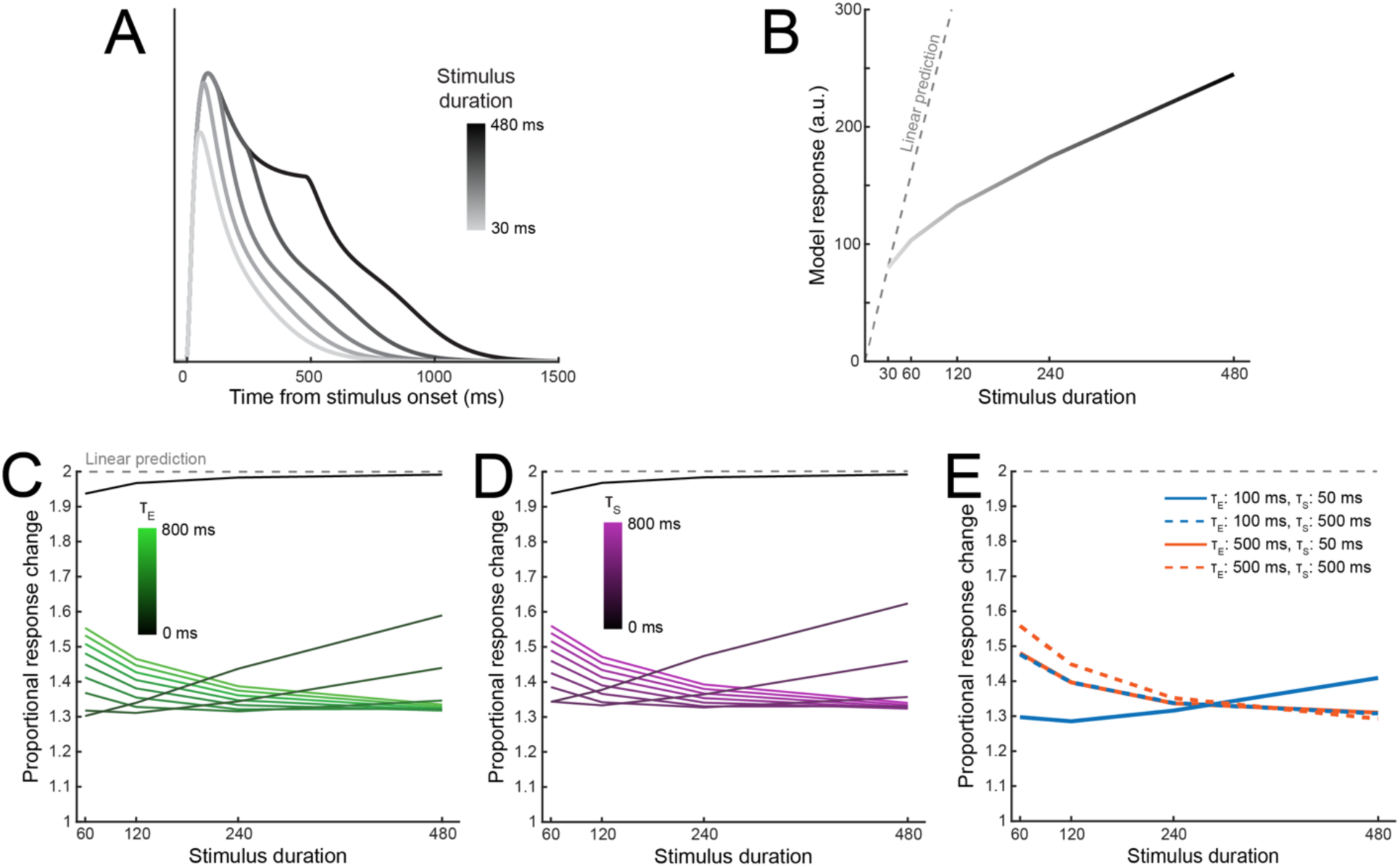
Subadditivity of model responses is robust across excitatory and suppressive time constants. A) Sensory layer responses over time in response to stimuli of varying durations (30, 60, 120, 240, or 480 ms), using τ_E_ = 100 ms and τ_S_ = 50 ms. B) Model responses calculated as the area under the curve of the sensory responses across the full simulated trial. Linear predictions are calculated relative to the model response for the 30 ms stimulus duration. C) Effect of τ_E_ and D) τ_S_ on subadditivity as a function of stimulus duration. Subadditivity was measured by taking the model response for each stimulus duration and dividing it by the response for a stimulus presented for half of that duration. Values below 2 represent subadditive responses. E) Proportional response changes as a function of different excitatory and suppressive time constants.

#### Response adaptation

Neurons exhibit response adaptation, such that responses are lower following repeated stimulus presentations (Kohn, 2007; Lisberger & Movshon, 1999; Priebe et al., 2002; Solomon & Kohn, 2014; Vogels, 2016). The magnitude of this adaptation depends on the interstimulus interval (ISI), with shorter ISIs resulting in stronger response adaptation. We observed a similar pattern in model simulations, where responses to the second stimulus (T2) in a sequence of two identical stimuli were lower when the ISI was shorter (e.g., 100 ms; Figure 6A) compared to longer ISIs (e.g., 900 ms; Figure 6B). We quantified the magnitude of response adaptation by measuring the reduction in the model response to T2 after subtracting out the response to T1 (i.e., the shaded blue region between curves in Figure 6A-B; (Lisberger & Movshon, 1999; Priebe et al., 2002). With increases in either τ_E_ or τ_S_, response adaptation persisted across longer ISIs (Figure 6C-D), as longer time constants allowed the excitatory and/or suppressive drives to extend across longer ISIs, leading to normalization of T2 by T1. For combinations of shorter and longer time constant parameters (Figure 6E), suppression accumulated as both τ_E_ and τ_S_ increased. Thus either excitatory temporal windows, suppressive temporal windows, or both could generate response adaptation, and had quantitatively similar effects on suppression indices.

**Figure 6.**
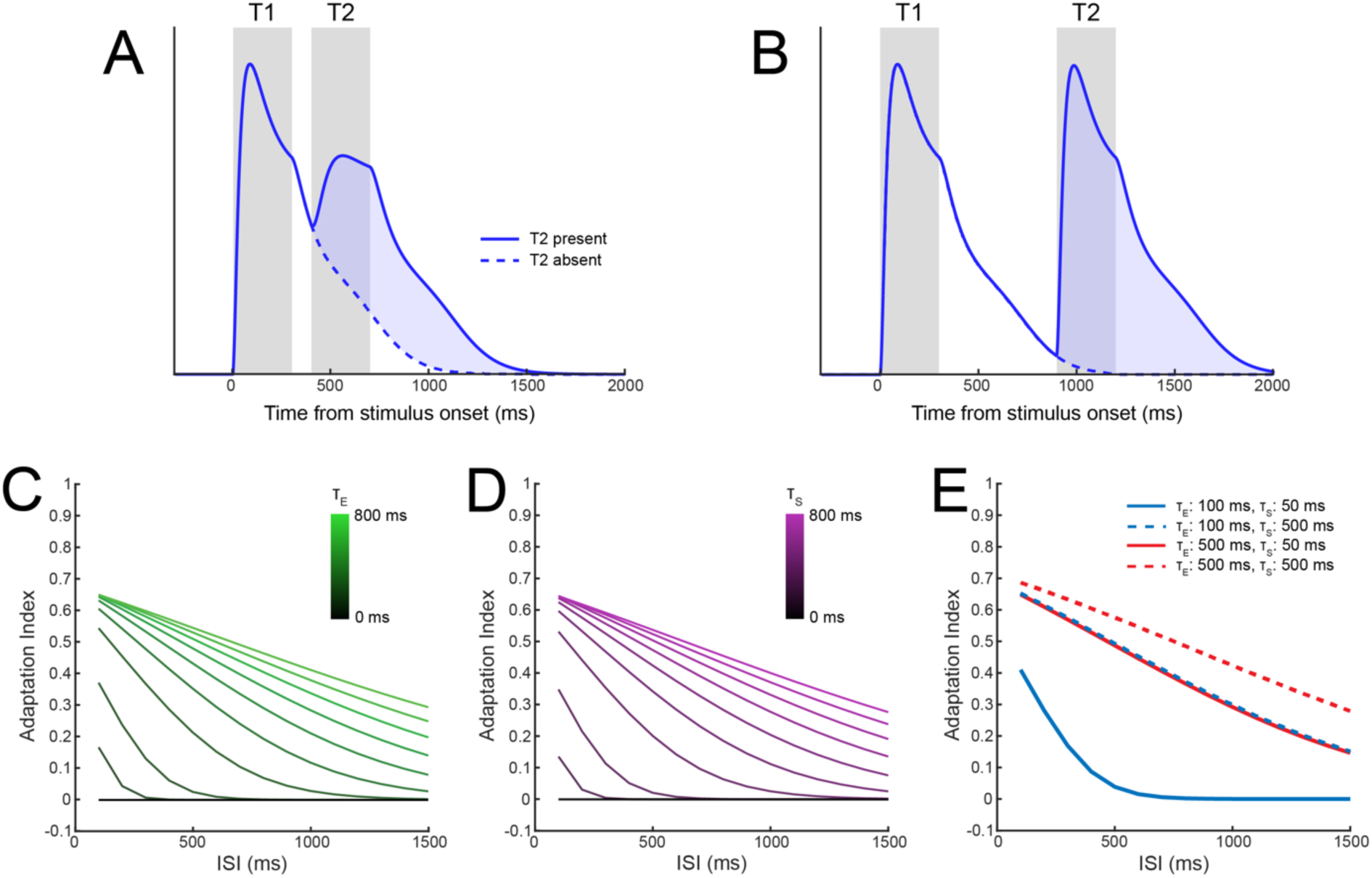
Response adaptation arises from either excitatory or suppressive temporal windows. A) Model sensory responses as a function of the presence of a preceding stimulus with an interstimulus interval of 100 ms, using τ_E_ = 100 ms and τ_S_ = 50 ms. Response adaptation is demonstrated by the reduced response to T2 (blue shaded region) relative to T1. B) Response adaptation was nearly eliminated when the ISI was increased to 600 ms. C) Effect of τ_E_ and D) τ_S_ on response adaptation for different ISIs. An adaptation index of zero represents no change in responses; positive indices show response adaptation. E) Combining different values of the τ_E_ and τ_S_ produces a range of response adaptation profiles.

### Spatiotemporal normalization captures phenomena not explained by spatial or temporal normalization alone

Whereas the above results could be achieved with existing models that implement either just spatial or just temporal normalization, several neural phenomena involve interactions between space (or features) and time. Here we tested whether D-STAN, with its unified spatiotemporal normalization computation, could capture such phenomena. For these purposes, features like orientation are treated equivalently to space, as in previous static normalization models (Reynolds & Heeger, 2009). The following analyses demonstrate how spatiotemporal normalization can generate suppression across stimuli with different feature values even when they are presented at different points in time. This property allows D-STAN to exhibit neural and behavioral effects found empirically but not seen in existing normalization models. As these next analyses show, a notable consequence of spatiotemporal normalization is that it leads to normalization-linked suppression both forward and backward in time.

#### Response adaptation by non-identical stimuli

Empirical work has shown that response adaptation can occur even with non-identical stimuli (Liu et al., 2009; Priebe & Lisberger, 2002), with one study in particular finding that most MT neurons are adapted by a wider range of motion directions than they are responsive to (Priebe & Lisberger, 2002). This is a crucial observation, as neurons that are suppressed only by their own past activity, as in DN models (Groen et al., 2022; Zhou et al., 2019), could not be suppressed by past stimuli that do not drive the neuron itself. In addition, neurons generally undergo less adaptation as the similarity between the adapting and test stimulus decreases (Liu et al., 2009; Patterson et al., 2013; Priebe et al., 2002). Therefore we measured model responses to a preferred target stimulus while varying the orientation of the adapting stimulus and calculated an adaptation index (see Methods; Figure 7A).

**Figure 7.**
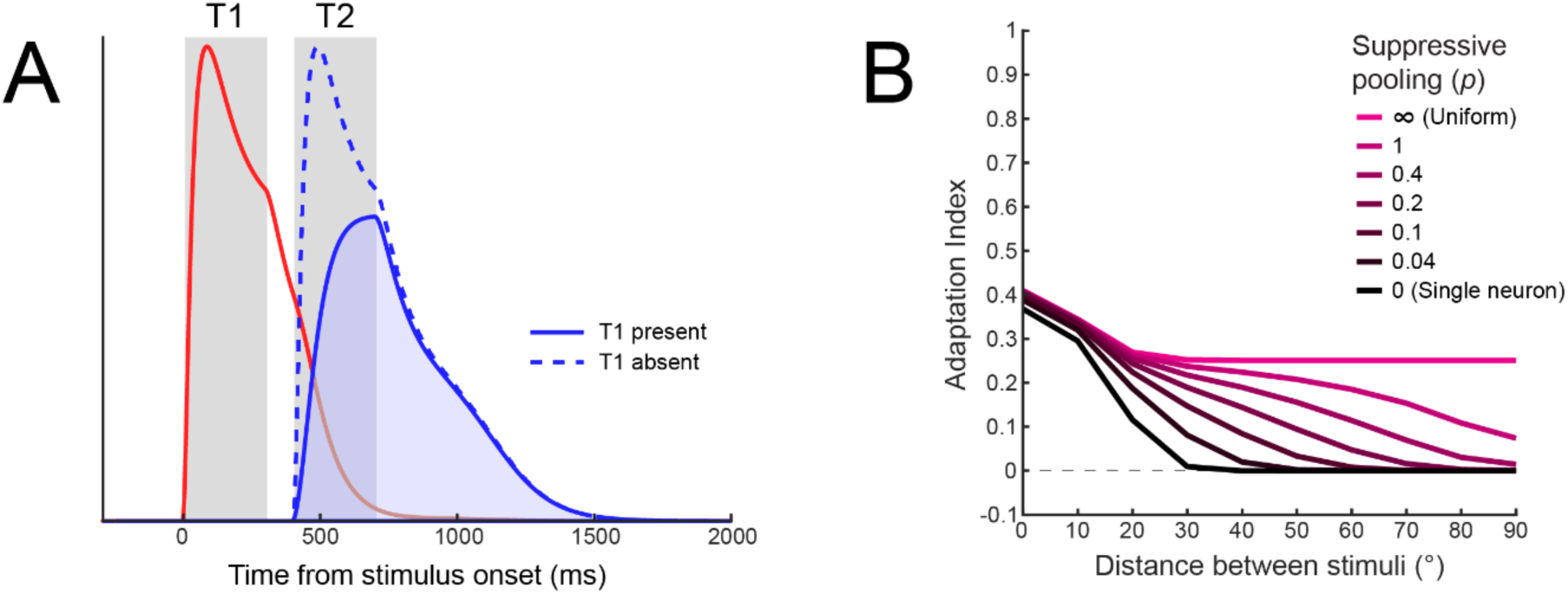
Response adaptation for non-identical stimuli in D-STAN. A) Model sensory responses as a function of the presence of a preceding stimulus with an orientation difference of 90° and interstimulus interval of 100 ms, using τ_E_ = 100 ms, τ_S_ = 50 ms, and uniform suppressive pooling. Response adaptation is demonstrated by the reduction in T2 responses while T1 is present (shaded blue vs. dotted blue region). B) Effect of tuned suppressive pooling on response adaptation for non-identical stimuli. When the suppressive drive was pooled uniformly, adaptation persisted over large orientation differences between the adapter and test stimuli. As the pooling tuning width decreased, adaptation decreased for more dissimilar stimuli.

We found, first, that adaptation was always strongest when the adapting stimulus and test stimulus matched in orientation. This property was due to the excitatory tuning of the model neurons: excitatory drives accumulated across both stimulus presentations when the neuron was sensitive to the adapting stimulus (<∼20° from the preferred orientation), resulting in stronger normalization of the test stimulus. Second, adaptation by non-identical stimuli depended on the tuning of the suppressive pool. When suppression was uniformly pooled across all sensory neurons (magenta line in Figure 7B), adaptation remained relatively stable across larger differences between adapting and test stimuli.

However, when suppression was adjusted to more strongly weight neurons tuned to similar orientations by changing the tuning width of the suppressive pool, adaptation progressively decreased and was even eliminated for more dissimilar orientations (Snow et al., 2016). In the limit, when a neuron is only suppressed by its own activity—as in DN models (Groen et al., 2022; Zhou et al., 2019)—adaptation occurs only for a narrow range of similar orientations. The ability to model different suppressive tuning profiles therefore allows D-STAN another way to capture effects that depend on more complex spatiotemporal interactions, compared to previous models.

#### Backward masking

In backward masking, a neuron’s ongoing response to an initial stimulus is suppressed by a subsequent stimulus (Breitmeyer & Öğmen, 2006; Enns & Di Lollo, 2000; Kovacs et al., 1995). Similar to response adaptation, the magnitude of backward masking diminishes as the stimulus onset asynchrony (SOA) between stimuli increases. D-STAN exhibited both backward masking and this characteristic SOA dependence, with greater masking at shorter SOAs (e.g., 250 ms; Figure 8A) than longer SOAs (e.g., 500 ms; Figure 8B). We simulated backwards masking using a sequence of two orthogonal stimuli and quantified the degree of masking in model simulations by measuring the response to T1 when T2 was present vs absent. Notably, the excitatory temporal window was necessary to produce backward masking, and its strength depended on τ_E_ (Figure 8C). Longer time constants resulted in backward masking that was generally stronger and persisted across longer intervals. In contrast, masking did not occur with the suppressive window alone, when τ_E_ was zero (Figure 8D). In this case, there was no temporal overlap in sensory responses to the two stimuli, which meant no normalization of the first stimulus by the second. However, when we explored the interaction between parameters, both τ_E_ and τ_S_ affected the magnitude of backward masking (Figure 8E). In general, higher τ_E_ values resulted in stronger backward masking, particularly for shorter SOAs (blue vs red lines in Figure 8E), as the extended sensory responses resulted in more overlap between the two stimuli and increased the overall normalization. In contrast, higher τ_S_ values resulted in reduced backward masking (dashed vs solid lines), as longer suppressive temporal windows tended to reduce the overlap in sensory responses between stimuli (cf. Figure 4D). Thus, the excitatory temporal window was necessary to produce backward masking, but so long as it was present, the strength of masking could be modulated by the suppressive temporal window.

**Figure 8.**
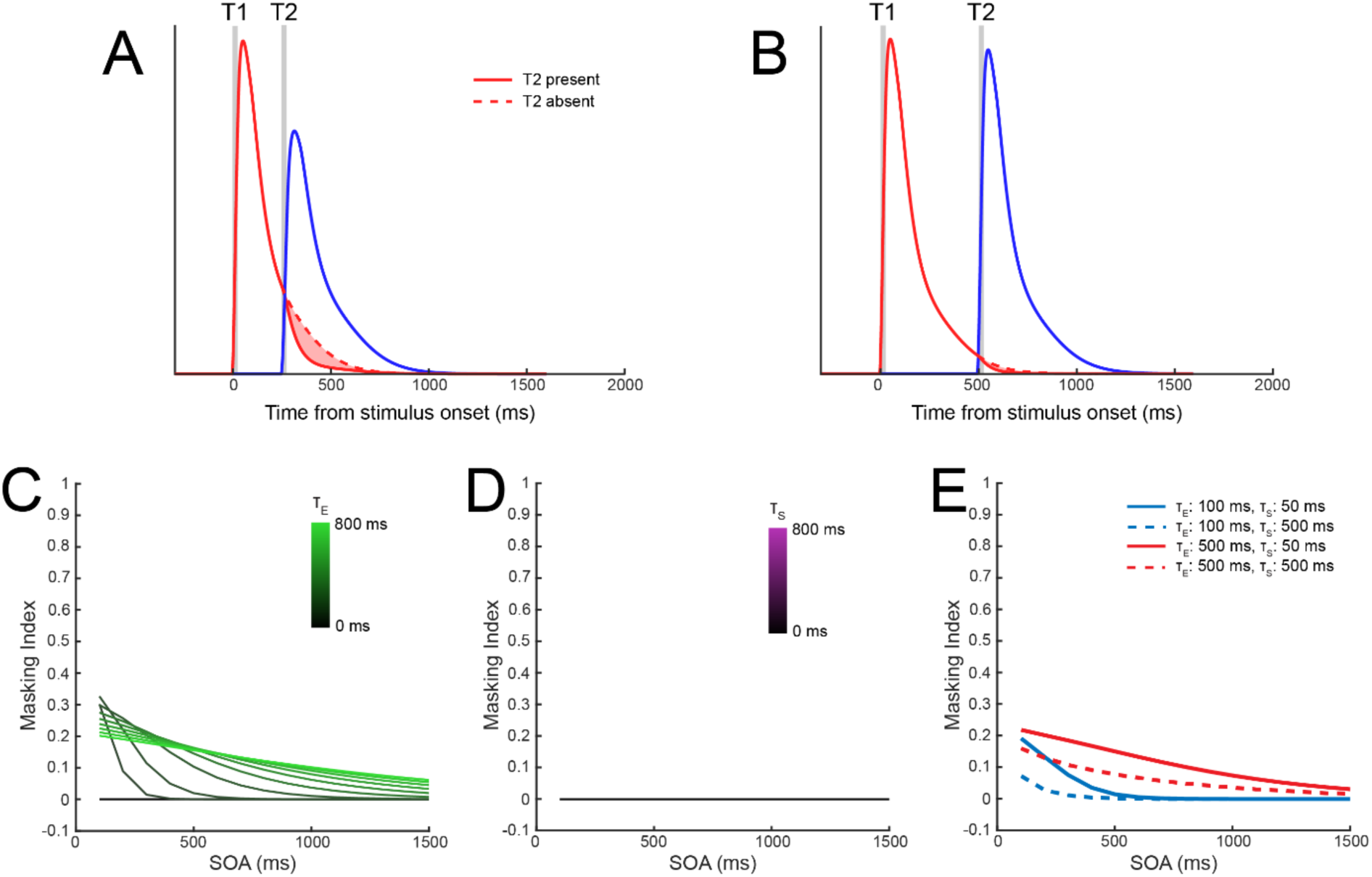
Backward masking requires excitatory temporal windows. A) Model sensory responses for one stimulus (T1, red lines) as a function of the presence of a subsequent stimulus (T2, blue line) with a stimulus onset asynchrony of 250 ms, using τ_E_ = 100 ms and τ_S_ = 50 ms. Backward masking is demonstrated by the reduction in T1 responses following the onset of T2, as indicated by the shaded red region. B) Backward masking was eliminated when the SOA was increased to 500 ms. C) Effect of τ_E_ and D) τ_S_ on backward masking for different SOAs. E) Combining different values of the τ_E_ and τ_S_ affected the pattern of backwards masking.

#### Contrast-dependent suppression from both past and future stimuli

A hallmark of spatial normalization models is that they generate suppressive effects that systematically depend on stimulus contrast (Heeger, 1992; Reynolds & Heeger, 2009). Therefore we next investigated the contrast dependence of temporal context effects produced by D-STAN. Specifically, we examined how model responses are affected by the contrast of a competing stimulus. In D-STAN, contrast increases the magnitude of stimulus input, and thus increases the overall excitatory and suppressive drives in the sensory layers. As such, we expected contrast modulations to have effects similar to our simulations of response adaptation and backward masking: higher contrast (presence vs absence) for one of two stimuli in a sequence would result in reduced model responses to the other.

We first simulated model responses using an SOA of 250 ms, with excitatory and suppressive time constants that resulted in both response adaptation and backward masking (τ_E_ = 400 ms, τ_S_ = 100 ms; see Figure 2C). When the contrast of the second stimulus (T2; Figure 9A) was high (64%) vs. low (16%) we observed a reduction in the model response to the first stimulus (T1). Likewise, when the contrast of T1 was high vs. low, the model response to T2 was reduced (Figure 9B). Both effects were due to increases in the overall suppressive drive caused by higher contrast stimuli, resulting in stronger normalization of the model response to the other stimulus.

**Figure 9.**
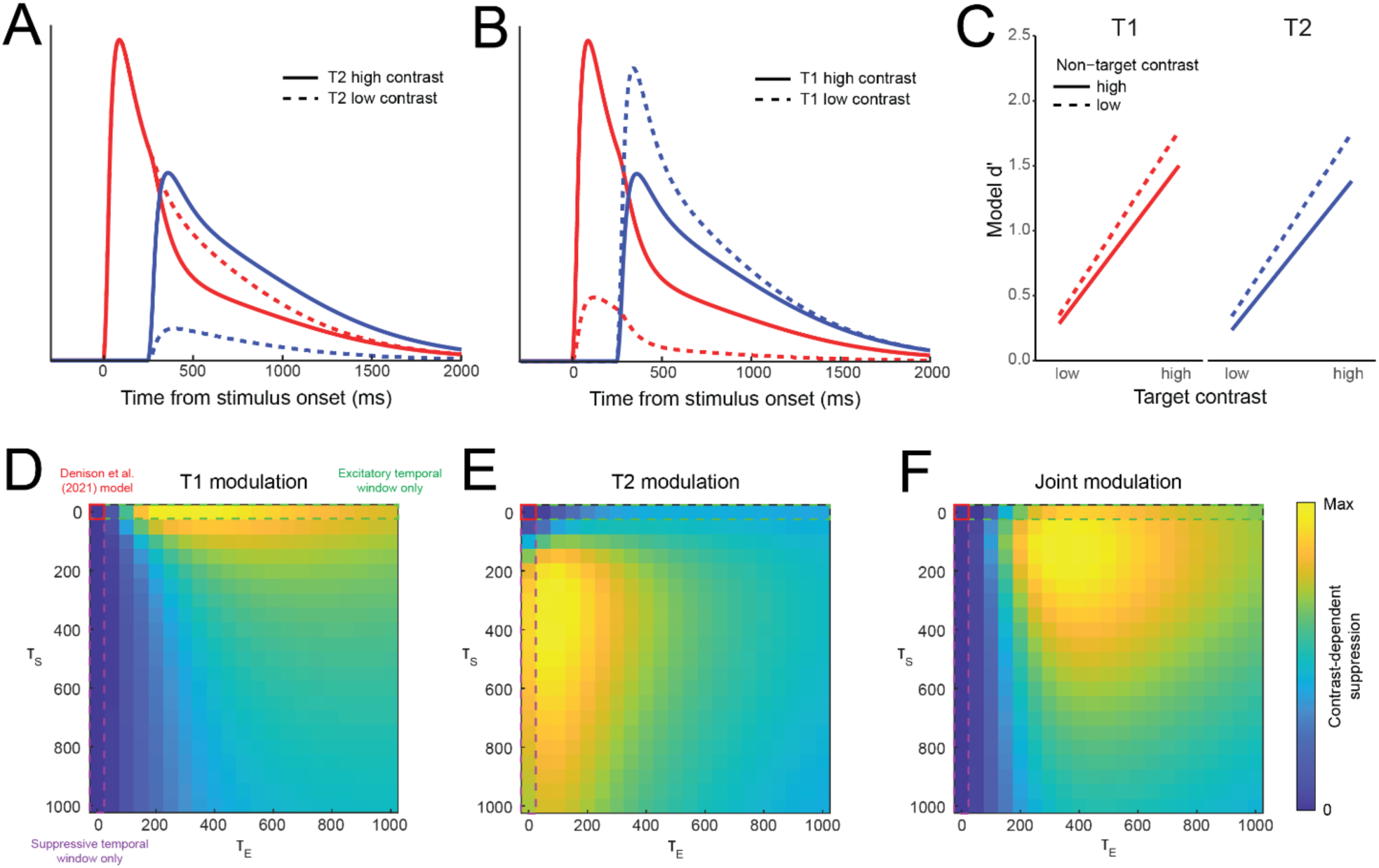
Contrast-dependent suppression forward and backward in time. A) When T1 (red lines) was fixed at high contrast, its response was modulated by the contrast of T2 (red solid line vs dashed line), using τ_E_ = 400 ms and τ_S_ = 100 ms. B) Likewise, the response to a high contrast T2 was modulated by the contrast of T1 (blue solid vs dashed lines). C) Modeled behavioral prediction in terms of d’ for discriminating clockwise vs counterclockwise stimulus orientations. Performance was lower for each target stimulus when the non-target was presented at higher contrast, consistent with recent empirical observations. D) Effect of τ_E_ and τ_S_ on the contrast-dependent modulation of T1, E) T2, and F) the joint modulation of both stimuli, calculated by a pointwise multiplication of individual target heatmaps. A region of the parameter space with moderate τ_E_ and low τ_S_ provided the strongest combined modulation of both target stimuli, comparable to human behavior as reported by Epstein and Denison (2023).

Psychophysical work has demonstrated that orientation discriminability for a target stimulus can be impaired by a non-target stimulus presented either before or after it by 250 ms, with greater impairment for higher vs. lower contrast non-target stimuli (Epstein & Denison, 2023). To determine if the model could also produce this behavioral pattern, we used the decision layer of D-STAN to calculate discriminability (d’) between different target orientations (Figure 9C). Model d’ was higher when the non-target was presented at a lower contrast, regardless of the target contrast. Notably, this effect occurred both forward (i.e., the contrast of T1 affected responses to T2) and backward in time (i.e., the contrast of T2 affected responses to T1). In these simulations, excitatory drives were affected by target contrast, with higher target contrast resulting in stronger sensory responses and higher model d’ for targets. On the other hand, the effects of non-target contrast were imparted through the suppressive drive, with high contrast non-targets resulting in lower sensory responses and d’ for targets.

We also explored how the excitatory and suppressive temporal windows modulated the effects of one stimulus’s contrast on the model responses to the other stimulus. We found that temporal windows were necessary for contrast-dependent suppression, as when both time constants were zero— reducing the model to the original Denison et al. (2021) version—there was no modulation for either stimulus (Figure 9D-F, red solid outline). For T1, we found that contrast-dependent suppression was strongest for a wide range of τ_E_ values (∼250-850 ms) combined with a low τ_S_ (0-100 ms; Figure 9D). In particular, we found that suppression alone (i.e., τ_E_ = 0) was insufficient to produce the observed effects (Figure 9D, purple dashed outline), because the lack of any extended excitatory drive meant the sensory responses did not overlap in time. This is comparable to the lack of backward masking present in our simulations manipulating only τ_S_ (cf. Figure 8D). In contrast, modulation of T2 was strongest when τ_S_ was an intermediate value (∼250-650 ms) and τ_E_ was low but non-zero (∼50-150 ms; Figure 9E). To assess what temporal window parameter combinations produced contrast-dependent suppression simultaneously across both T1 and T2 (Figure 9F), we computed a combined modulation index, which showed a range of intermediate τ_E_ values (300-600 ms) combined with low τ_S_ (50-200 ms) that resulted in moderate suppressive effects for both T1 and T2 (as in Figure 9C, with τ_E_ = 400 ms and τ_S_ = 100 ms). The ability of D-STAN to produce bidirectional contrast-dependent suppression demonstrates how temporal normalization can account for interactions between stimuli that are separated in time, both affecting ongoing sensory processing as well as behavioral responses.

## Discussion

Normalization is a successful computational principle that predicts neural responses and perception, yet most previous models have considered normalization across space or time alone. Here, we demonstrate how spatiotemporal normalization can be implemented in a neural network model of dynamic visual processing. D-STAN is grounded in the framework of normalization established by Reynolds and Heeger (2009), which was subsequently extended into a dynamic spatial normalization model by Denison et al. (2021). The core of D-STAN is a unified spatiotemporal normalization computation, which provides local contextual modulation across space, time, and features. This normalization computation is supported by spatiotemporal receptive fields, with excitatory and suppressive drives that depend on recent stimulus history. This architecture generalizes across three classes of previous models: 1) static spatial normalization models (Reynolds & Heeger, 2009), which lack a temporal dimension; 2) delayed normalization models (Groen et al., 2022; Zhou et al., 2019), which include temporal normalization but lack spatial and featural dimensions, and; 3) the normalization model of dynamic attention (Denison et al., 2021), a dynamic model which includes normalization across space and features, but lacks temporal normalization. Spatiotemporal normalization in D-STAN encompasses all these previous models, while also going beyond their simple combination, allowing D-STAN to generate additional phenomena not previously captured.

We report several findings. First, the model sensory neurons exhibit temporal receptive fields with biphasic profiles, similar to neuronal receptive field properties observed empirically (Mante et al., 2008). Notably, this biphasic profile was not incorporated directly into the model but emerged from the interaction between the excitatory and suppressive drives, the exponential temporal windows, and the normalization computation. Second, D-STAN exhibited surround suppression, consistent with static spatial normalization models (Carandini & Heeger, 2012); and it reproduced several non-linear dynamic response properties, including transient-sustained response dynamics (Lisberger & Movshon, 1999; Motter, 2006), subadditivity with increasing stimulus duration (Groen et al., 2022; Zhou et al., 2018), and response adaptation/repetition suppression (Kohn, 2007; Priebe et al., 2002; Vogels, 2016), consistent with previous delayed normalization models (Groen et al., 2022; Zhou et al., 2019). Third, D-STAN goes beyond both spatial and delayed normalization models to predict a range of phenomena that depend on spatiotemporal or feature-temporal interactions: adaptation by non-identical stimuli (Priebe & Lisberger, 2002), backward masking (Breitmeyer & Öğmen, 2006; Enns & Di Lollo, 2000; Kovacs et al., 1995), and bidirectional contrast-dependent suppression between successive stimuli, which has recently been observed psychophysically (Epstein & Denison, 2023). D-STAN can thus account for a wide range of neural and behavioral findings in the domain of dynamic vision through spatiotemporal normalization.

We implemented spatiotemporal normalization in D-STAN by allowing the excitatory and suppressive drives to depend on recent stimulus input via their “temporal windows”, the temporal component of their spatiotemporal receptive field structure. These temporal windows cause the excitatory and suppressive drives to persist within the model sensory layers even after a stimulus has offset, allowing for normalization to occur between stimuli that are presented at distinct points in time. We found that the excitatory and suppressive temporal windows contributed to the observed phenomena in different ways. Increasing τ_E_ carried neural responses forward in time, after stimulus input had ended (Figure 4A), such that these responses could suppress (and be suppressed by) responses to subsequent stimuli. This was necessary for reproducing suppression backward in time (i.e., backward masking or contrast-dependent suppression from a subsequent stimulus), because responses needed to be carried forward across interstimulus intervals so that normalization could occur between neurons tuned to different features. In contrast, increasing τ_S_ allowed suppressive drives to accumulate across longer intervals, increasing the overall suppression of responses to prolonged stimulus presentations, resulting in transient-sustained responses typical of neural firing (Figure 4D), but also enabling suppression to be carried forward after responses to an initial stimulus have ended. This extended suppressive drive was sufficient to reproduce suppressive effects forward in time (i.e., response adaptation and contrast-dependent suppression from a preceding stimulus). For phenomena isolated within a single neuron (e.g., subadditivity, response adaptation by an identical stimulus) the excitatory temporal window reproduced equivalent patterns since history-based changes in the excitatory drive propagate to the suppressive drive, even when suppression is only instantaneous (τ_S_ = 0 ms). Thus, the excitatory and suppressive temporal windows had distinct effects on neural timecourses, resulting in different patterns of effects across the phenomena we investigated. Importantly, combining both the excitatory and suppressive temporal windows in D-STAN was necessary to produce this wide array of phenomena.

Recent models have attempted to account for several of the dynamic neural response properties we investigated here. Delayed normalization (DN) models, for example, divide a neuron’s excitatory drive by a filtered and delayed copy of itself, generating neural response time courses that can be fit to observed fMRI (Zhou et al., 2018) or electrocorticography data (Groen et al., 2022; Zhou et al., 2019), producing transient-sustained responses, subadditivity, and response adaptation. Compressive spatiotemporal (CST) models fit separate sustained and transient channels that are passed through a compressive nonlinearity to produce responses (Kim et al., 2024; Kupers et al., 2024). Although CST models have not been tested for the same set of response dynamics, they are well fit to fMRI BOLD time courses in response to sequences of stimuli with different spatial locations and timings, and they reproduce increasing temporal window sizes along the visual hierarchy (Murray et al., 2014; Wolff et al., 2022). In both types of models, normalization is computed independently within each recorded unit (voxel, electrode site, single-unit, etc.). In contrast, D-STAN implements suppressive pooling, both spatially—via summation across model neurons within layers—and temporally—as a consequence of the temporal windows—such that normalization can be induced by activity generated by other units and at other times. Pooling suppression across neurons is a property key to previous foundational normalization models (Reynolds & Heeger, 2009) and allows for stimuli outside of the “classical receptive field” (i.e., stimuli that do not by themselves drive a neuron) to suppress a neuron’s response. Spatial pooling allows D-STAN to capture phenomena that are not necessarily localized to a single unit, such as backward masking, contrast-dependent suppression, and response adaptation to non-identical stimuli.

Additionally, while DN and CST models both produce continuous neural time courses, these are computed based on *a priori* knowledge of the full stimulus sequence. D-STAN, in contrast, produces layer responses in a recursive manner, with computations implemented “online” at each timestep.

Other models have implemented normalization in a dynamic framework. Louie et al. (2014) modeled decision circuits in LIP using excitatory and inhibitory model neurons, noting that the recurrent interaction between neurons resulted in normalization that depended on an exponential weighting of previous excitatory activity. The suppressive temporal window in D-STAN matches with this formulation, demonstrating that temporal normalization can explain sensory phenomena, as well as the dynamics of decision making. More recently, Ernst et al. (2021) used a model similar to that of Louie et al. (2014) to predict a variety of transient-sustained responses recorded from MT neurons. Their dynamic equations used separate time constants for updating the excitatory and inhibitory responses over time, and notably in almost all neurons the fitted inhibitory time constant was longer than excitatory time constants. In D-STAN, this property—slower suppression than excitation—is a necessary consequence the model structure, since suppressive drives depend on both τ_E_ and τ_S_, so even when τ_S_ is shorter than τ_E_, the effective time constant of suppression is longer than τ_E_. It remains to be determined whether this is indeed a universal property in cortex, as predicted by D-STAN.

Spatial normalization across local neuronal populations has been proposed as a canonical computation in the brain that may provide benefits to coding efficiency (Carandini & Heeger, 2012; Louie & Glimcher, 2012), and evidence for it has been found in a variety of cortical regions (Busse et al., 2009; Carandini et al., 1997; Louie et al., 2011; Rabinowitz et al., 2011; Simoncelli & Heeger, 1998; Zoccolan et al., 2005). In a similar way, temporal normalization may enhance sensitivity to changes in the environment, with transient-sustained dynamics, subadditive neural responses, and response adaptation demonstrating that consistent inputs act to reduce overall neural activity (Zhou et al., 2019; Zhou et al., 2018). Temporal normalization may also emphasize differences between stimuli across time, as in the contrast-dependent suppression effects we observed, where model d’ was increased for high contrast targets among low contrast non-targets. Notably, previous formulations of temporal normalization capture forward suppression (e.g., adaptation) but not backward suppression, but coding efficiency should benefit from reducing redundancies bidirectionally in time. D-STAN provides such a bidirectional normalization mechanism. These effects on coding efficiency also depend on how suppression is pooled across a neural population, as we observed when we manipulated suppressive tuning width during response adaptation: if suppression more strongly weighs inputs from similarly tuned neurons (i.e., larger *p* in Figure 7B), it can increase sensitivity to larger changes in stimulus features over time. Conversely, broader tuning offers robustness against small perturbations caused by noise and can promote stability of representations over time (Benucci et al., 2013; Snow et al., 2016).

D-STAN is a flexible model that can be extended in different ways. First, the model can handle different spatial and feature dimensions through the tuning properties of the sensory neurons. In the current study, we manipulated the spatial tuning of sensory layer neurons to examine surround suppression and the tuning of the suppressive pool to investigate feature-tuned adaptation. Previous DN models, in which suppression was calculated separately for each neural unit (Groen et al., 2022; Zhou et al., 2019), are essentially a special case of tuned suppression, where the suppressive pool is tuned narrowly to one neuron. Second, we defined the excitatory and suppressive temporal windows using exponential functions, but different parameterizations are possible. Third, it may be interesting to examine how the modeled excitatory and suppressive time constants relate to the dynamics of excitatory and inhibitory neurons within different neural circuits. Recent computational work has demonstrated how normalization can be implemented through recurrent excitation and inhibition (Heeger & Zemlianova, 2020), providing a new class of models with which to examine dynamic processing at the level of cortical circuits. Fourth, D-STAN can be expanded to include multiple hierarchical sensory layers with different spatiotemporal receptive fields and pooling profiles. Neural dynamics vary between different regions, and temporal integration windows have been found to increase along the cortical hierarchy (Chaudhuri et al., 2015; Fritsche et al., 2020; Gao et al., 2020; Murray et al., 2014; Vidaurre et al., 2017; Wolff et al., 2022), suggesting that temporal windows may be successively applied and accumulate over several stages of processing. Finally, D-STAN has the capacity to model the dynamics of voluntary and involuntary attention—which were not included in the current simulations—and so a natural extension is to examine how attention interacts with spatiotemporal normalization to influence ongoing processing.

D-STAN makes several testable predictions for future empirical work. 1) Single-neuron responses will be well-fit by D-STAN, and the excitatory and suppressive spatiotemporal tuning parameters fit using one stimulation protocol (e.g., reverse correlation) will allow quantitative predictions of how that neuron will behave during other stimulation protocols that elicit a range of spatiotemporal phenomena: subadditivity, response adaptation, backward masking, etc. 2) Biphasic temporal receptive fields arise not from subtractive inhibition but rather from divisive suppression of past time points. As a result, they will only be revealed when temporal receptive fields are mapped using continuous visual stimulation, as during reverse correlation protocols, and not using single stimulus presentations. Such techniques have been used to map spatiotemporal receptive fields in mammalian LGN (Cai et al., 1997; Mante et al., 2008), V1 (De Valois et al., 2000), and MT (Perge et al., 2005), as well as in visual regions in Drosophila (Sun et al., 2017). While spatiotemporal receptive fields have mostly been investigated for their relevance to motion perception, our findings suggest that they may play a much broader role in visual processing. 3) Suppressive fields governing normalization will be not only spatially broader and temporally more extended, but *spatiotemporally* broader than their excitatory field counterparts, predicting that neurons will exhibit adaptation to stimuli that fall outside their classical receptive fields. Initial observations supporting this prediction come from Priebe et al. (2002), but more experiments are needed to test the generality of this prediction and to characterize the spatiotemporal structure of suppressive fields. An open question amenable to empirical measurements would be whether spatial and temporal suppressive field components are independent—with the spatial tuning of suppression constant across time, as we have modeled here—or interacting, such that the spatial tuning of suppression changes (likely, narrows) further into the past.

In summary, we introduced the dynamic spatiotemporal attention and normalization model, D-STAN. The model integrates standard spatial and feature-based neural tuning functions with excitatory and suppressive temporal windows to generate a unified spatiotemporal normalization computation, resulting in a spatiotemporal receptive field structure like that seen in physiological recordings of sensory neurons. D-STAN reproduces several non-linear response properties, including subadditive responses, response adaptation, and backwards masking, as well as contrast-dependent suppression between stimuli across time, a perceptual phenomenon only recently shown in human observers (Epstein & Denison, 2023). Our model provides advances over other dynamic normalization (Denison et al., 2021; Groen et al., 2022; Zhou et al., 2019) or compressive spatiotemporal (Kim et al., 2024; Kupers et al., 2024) models, such as the recursive neural network architecture that allows for continuous online prediction of layer responses, as well as the flexibility afforded by the temporal window and suppressive pooling structures, which allows both spatial and temporal normalization to be carried out via a single, parsimonious spatiotemporal computation. Overall, D-STAN provides an important step toward the goal of developing real-time process models of dynamic vision.

## Materials and Methods

### Code Accessibility

Model code and analysis scripts are available on the project Github repository: https://github.com/denisonlab/dstan. Data produced by several of the analyses are available on OSF: https://osf.io/qy9pa/.

### Model specification

D-STAN is a hierarchical, recurrent neural network, which models the dynamics of feature-tuned neural populations given time-varying sensory input. D-STAN consists of interconnected sensory and decision layers, that each produce time-varying responses in model neurons, allowing the model to generate predictions about neural activity from continuous input in an online fashion (i.e., time step by time step) as well as to generate predictions about behavioral performance in simple perceptual tasks (Figure 1). D-STAN is built on a modeling foundation established by Denison et al. (2021) and Reynolds and Heeger (2009). The introduction of excitatory and suppressive temporal windows together with a spatiotemporal normalization computation allows D-STAN to generate a rich repertoire of dynamic behavior and to exhibit effects of temporal context in line with empirical observations, which could not be generated by these previous models. D-STAN also contains voluntary and involuntary attention layers, following Denison et al. (2021), though these were removed for all analyses in the current study.

### Sensory layer

The sensory layer represents the visual processing stage of the model. As in the normalization model of dynamic attention (Denison et al., 2021), this layer receives stimulus input at each time point, which feeds into N = 12 neurons that are tuned to different feature values. We use orientation as an example feature throughout the reported analyses. In simulations, the time course of stimulus input was represented in the matrix ***X*** (with size of M orientations x K time points), with each vector 𝑿_𝒕_= (0, 0, …, *c*, …, 0) indicating the currently presented orientation at contrast level *c*. When no stimulus was presented, all elements of the vector were zero. The stimulus drive to each neuron at each time point depended on the match between the stimulus orientation and the neuron’s orientation tuning function, as determined using a raised cosine function:

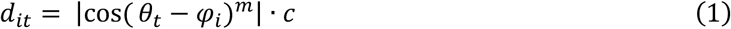

where 𝜃_𝑡_is the orientation of the stimulus shown at time *t*, 𝜑_𝑖_ is the preferred orientation of the *i*th neuron, and *c* is the stimulus contrast. Orientation tuning functions were evenly spaced across the feature space, 𝜑_𝑖_ = π(𝑖 − 1)/N. The exponent *m* controls the width of the tuning curve and was set to 23 (*m* = 2N – 1).

Unlike in previous models, the excitatory drive for each neuron is calculated at each time point using a weighted exponential of recent stimulus drive:

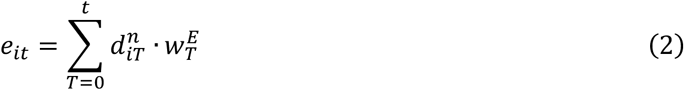

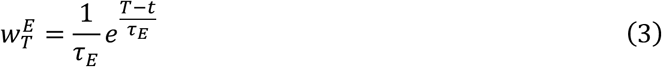

where *t* is the current time point, 𝑛 is an exponent that affects the shape of neurons’ contrast response functions, and 𝜏_𝐸_ is the time constant determining the amount of weight given to previous time points. In effect, this excitatory temporal window imbues the model neurons with a temporal receptive field, where not only are units responsive to their preferred stimulus at any given time point, but also respond based on the stimulus history as well.

The suppressive drive was also calculated at each time point, and weighted by an exponential:

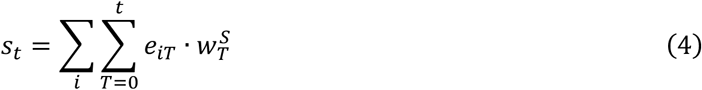

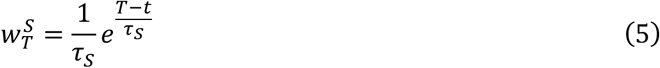

Thus, a neuron’s current response is suppressed by the activity history of itself and its neighbors, which together comprise the suppressive pool. In the reported simulations, only a single spatial location was modeled, and neurons tuned to all orientations were included with equal weight in the suppressive pool. But in principle, the suppressive pool could also be tuned to locations and features.

Finally, the response of each neuron was updated at every time step using the following differential equation, which functions as a dynamic update of the standard normalization equation (Reynolds & Heeger, 2009), as in the normalization model of dynamic attention (Denison et al., 2021):

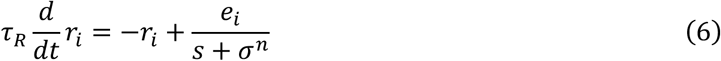

where 𝑟_𝑖_is the response of the *i*th neuron within the sensory population, 𝜏_𝑅_ is a time constant that affects the rate at which the neuron’s response increases during stimulus presentation and decreases after stimulus offset, 𝑒_𝑖_ is the excitatory drive of that neuron, 𝑠 is the (pooled) suppressive drive, 𝜎 is a semi-saturation constant that affects the contrast gain of neurons and keeps the denominator non-zero, and 𝑛 is an exponent following Eq. 2. Some parameter values (𝜏_𝑅_ = 52, 𝑛 = 1.5) were fixed as fitted to empirical data in previous reports (Denison et al., 2021), however we decreased the semi-saturation constant (𝜎 = 0.1) to account for the fact that we only used a single sensory layer in this version of D-STAN.

### Decision layer

The decision layer receives input from the sensory layer, which it uses to compute behavioral output regarding the orientation of the stimuli, as in the normalization model of dynamic attention (Denison et al., 2021). Decisions are generated by two neurons, each encoding for one of the two stimuli (T1 and T2). The excitatory drive for each decision neuron is:

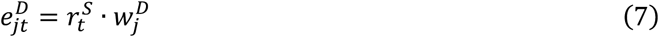

where each vector 𝑤^𝐷^is designed to compute an optimal linear readout from the sensory responses to the *j*th stimulus, to determine whether the stimulus was rotated clockwise vs. counterclockwise from horizontal or vertical. The vectors project the sensory layer response onto the difference between two templates encoding the population responses to the CW and CCW stimuli along a given orientation axis; the axis of each stimulus (vertical or horizontal) is assumed known. Evidence accumulation is positive for CW decisions, and negative for CCW decisions, such that the sign of the evidence indicates the decoded orientation within the decision layer, while the magnitude of the evidence indicates the strength of the model decision. The suppressive drive is pooled over the decision neurons, and responses are updated at each time point according to the same differential equation as in sensory layers (𝜏_𝐷_= 10^5^, 𝜎 = 0.7; (Denison et al., 2021). The long time constant of this layer allows for sustained evidence accumulation.

In contrast to previous model iterations (Denison et al., 2021), each neuron in the decision layer accumulates evidence throughout the duration of the simulated trials, rather than during discrete windows following each stimulus presentation. Because the effects of temporal normalization occur when the excitatory and suppressive drives elicited by the two stimuli overlap, the discrete decision windows were unable to capture any behavioral effects of normalization for T1 at short stimulus onset asynchronies (SOAs). At the end of each simulation, the accumulated evidence for each stimulus is converted into *d’*, a measure of perceptual sensitivity, through multiplicative scaling (s_T1_ = s_T2_ = 1×10^5^).

### Simulation procedures

All simulations were performed in MATLAB (2022b, MathWorks, Natick, MA). We used a time step Δ𝑡 of 2 ms throughout simulations. The duration of the simulated trial was varied as necessary to capture model dynamics. When a stimulus was presented, we used a standard contrast level of 64% and presentation duration of 30 ms, unless otherwise specified. We varied the two temporal window time constants 𝜏_𝐸_ and 𝜏_𝑆_across simulations to assess how the temporal windows affected sensory layer dynamics and decision layer outputs.

### Estimating temporal receptive fields of model neurons

To assess how stimulus input at different past time points affects model responses at the current time, we performed a simulation and analysis based on reverse correlation, similar to the way a temporal receptive field might be measured in a neurophysiology experiment (Mante, 2008). We modified the stimulus input to the model to be a random binary vector, such that the stimulus drive at each time point was one or zero. We then performed model simulations for 10,000 different random stimulus vectors with a duration of 1200 ms, resulting in variable excitatory and suppressive drives and sensory layer responses that depended on the stimulus history. To estimate the impact of the stimulus input on each model timeseries, we used reverse correlation to calculate the average change in each measure caused by the presence of a stimulus at each time point by correlating the stimulus input at each time point with the response (excitatory/suppressive drive or layer response) at the final time point (Ringach & Shapley, 2004). Shufled weights were calculated in the same manner after rearranging the stimulus input vectors so that they were not aligned with the responses from the same simulation.

To characterize the temporal receptive fields and enable comparisons across different parameter settings, we fit the estimated stimulus weights using different functional forms. For the excitatory drive, we compared the estimated stimulus weights with the exponential function that defines the excitatory temporal window. This function (𝜏_𝐸_ = 400 ms) had no free parameters, but was adjusted using a scaling parameter to align it to the magnitude of the stimulus weights (the units of which are arbitrary), estimated using simple linear regression. For the suppressive drive, we similarly overlaid a scaled function to the stimulus weights. This function, *h_S_*, was determined by convolving the excitatory and suppressive temporal windows, using their respective time constants. We zero-padded the temporal windows at positive (i.e., future) time points to achieve the resulting functional form. Notably, this

function in general can also be calculated by taking the difference between the exponential temporal windows (for t < 0 and 𝜏_𝐸_ ≠ 𝜏_𝑆_):

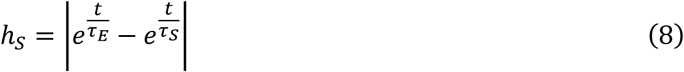

The sensory layer response weights were fitted with a difference of Gamma functions, using the simplified Gamma function form adopted in previous work (Zhou et al., 2019):

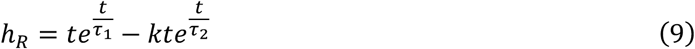

with time constants 𝜏_1_and 𝜏_2_, and weight 𝑘. We fitted the simulated stimulus weights to this function in MATLAB by minimizing the least-squares error using *fminsearch* up to a scaling factor. We performed this optimization 100 times, with random initial values for 𝜏_1_ and 𝜏_2_ drawn uniformly between 0 and 900, and initial *k* = 0, and selected the best fitting function across all solutions. For the function shown in Figure 2C, the best fitting parameters were: 𝜏L_1_ = 305.01, 𝜏L_2_ = 61.98, 𝑘^P^ = 5.43.

To assess how the effective temporal receptive fields were affected by the excitatory and suppressive temporal parameters, we performed the model simulations again for each combination of 𝜏_𝐸_ and 𝜏_𝑆_from 100 to 900 ms in 100 ms steps. In the results shown in Figure 2D-E, we took the fitted functions for one parameter fixed at 400 ms, while the other parameter varied.

### Surround suppression

To confirm that D-STAN maintains the spatial normalization properties observed in previous static models, we simulated how the model response to a stimulus was affected by a nearby stimulus presented outside its receptive field. To model the spatial dimension, we defined two populations of neurons each with feature tuning to multiple orientations, but tuned to distinct spatial locations. Whereas each subpopulation was excited only by stimuli in its preferred spatial location, suppressive drives were pooled across the two subpopulations (i.e., across space), allowing for mutually suppressive interactions between the two spatial locations. We assigned one subpopulation as “center” neurons, and the other as “surround” neurons. Across simulations, we varied the contrast of the center stimulus across a wide range of values (from 5% to 100% contrast at 20 levels, log-spaced) to map the contrast response function (CRF) of model neurons, and varied the contrast of the surround stimulus at fixed levels (0%, 12%, 25%, 50%, or 100%) to assess surround suppression. For each simulation, we measured the response in the neuron maximally responsive to the center orientation, summed across the timecourse of the simulation. We used a fixed set of time constants (τ_E_= 400 ms, τ_S_ = 100 ms), presenting the center and surround stimuli simultaneously for 100 ms. We also used a lower semi-saturation constant for this simulation (𝜎 = 0.02) to produce CRFs that spanned the neuron’s response range. The implementation of just two adjacent spatial populations is a simplification compared to previous implementations of spatial normalization that model a continuous spatial map (Carandini, 2004; Reynolds & Heeger, 2009). Nevertheless, this simplified implantation is sufficient to test whether the model can reproduce spatial normalization effects such as surround suppression, and it demonstrates how the proposed architecture can be flexibly expanded as needed to generate spatial and feature maps in the sensory layer.

### Effects of temporal receptive fields on neural responses

To assess the effects of the excitatory and suppressive time constants on the model neuron responses, we simulated trials in which a single target stimulus was presented. The stimulus presentation duration was set to 2000 ms to allow time to observe the peak and steady state responses, and the total trial duration was set to 8100 ms, including a 500 ms prestimulus period. We varied 𝜏_𝐸_ and 𝜏_𝑆_separately across 9 levels (50, 100, 200, 300, 400, 500, 600, 700, and 800 ms). To examine the effects of the individual time constants in isolation, for simulations varying 𝜏_𝐸_, we fixed 𝜏_𝑆_at zero, and vice-versa, effectively removing the excitatory or suppressive temporal receptive fields from the model. Setting 𝜏_𝐸_ = 𝜏_𝑆_ = 0 reduces the model to the previous version of Denison et al. (2021). For these simulations, we extracted response time courses from the sensory layer neuron tuned nearest to the target orientation. We also examined the interaction of the time constants by conducting simulations with different combinations of non-zero values for 𝜏_𝐸_and 𝜏_𝑆_.

For simulations varying the excitatory time constant, all model neurons reached the same stable activity level, and all neural responses were scaled by dividing the response at all time points by the maximum response within the stimulus presentation window. For each value of 𝜏_𝐸_, we found the time point at which the model responses reached 99% of the maximum response (“Time to peak”, Figure 3B), and the time point at which responses fell to 50% of the peak value following stimulus offset (“Time to half maximum”, Figure 3C). We used 99% of the maximum for the time to peak analysis, because responses approached but did not necessarily reach the numerical maximum until late in the window.

For simulations varying the suppressive time constant, we scaled all responses relative to the first peak (Figure 3D). Shorter suppressive time constants typically resulted in a faster reduction in model responses, reaching a peak sooner but also reducing the overall magnitude. To examine response dynamics relative to the peak response, we therefore normalized the response time course by the peak response during the stimulus presentation window. We calculated the time to peak by measuring the time at which the maximum response was reached (“Time to peak”, Figure 3E), as well as the model response 2000 ms after stimulus onset relative to the peak (“Stable activity level”, Figure 3F).

### Subadditive responses

To assess how neural responses depended on stimulus presentation duration, we again simulated trials with only a single target. For the simulations shown in Figure 5A, we used short time constants (τ_E_ = 100 ms, τ_S_ = 50 ms) and varied the stimulus duration by doubling from 30-480 ms, for 5 total duration conditions (30, 60, 120, 240, and 480 ms). We extracted responses from the sensory layer in the model neuron tuned closest to the target orientation, and calculated the model response by summing the sensory response across the entire trial duration, approximating the area under the curves in Figure 5A. To quantify subadditivity in neural responses, we calculated the effect of doubling the stimulus duration on the model response by dividing the response at duration 2x by the response at duration x (e.g., we divided the model response at 60 ms by that at 30 ms; Figure 5C). To assess how subadditivity depends on the excitatory and suppressive time constants, we selected one shorter and one longer value (𝜏_𝐸_ = 100 or 500 ms, 𝜏_𝑆_ = 50 or 500 ms), and computed the proportional response change for each pair of time constants (Figure 5D).

### Response adaptation

Response adaptation refers to the finding that neural responses to stimuli presented shortly after an initial stimulus are typically reduced in magnitude (Kohn, 2007; Motter, 2006; Priebe et al., 2002; Solomon & Kohn, 2014; Vogels, 2016). In our analysis, we therefore aimed to examine how model responses differed to a sequence of two “target” stimuli (referred to as T1 and T2) relative to a single stimulus. We performed separate analyses where T1 and T2 were identical stimuli (i.e., the same feature) or distinct (i.e., orthogonal features). In the first simulation examining the interaction between responses to two identical stimuli, our goal was to assess the magnitude of the response to T2 while subtracting out the activity related to T1. Therefore, we first measured the response to T1 alone (Figure 6A, blue dashed line), and then quantified the activity related to T2 by calculating the difference in responses when T2 was present vs. absent (Figure 6A, blue shaded region). We used a longer stimulus presentation duration of 300 ms, which is typical in the response adaptation literature (Vogels, 2016). For the simulation shown in Figure 6A-B, we again selected short time constants (τ_E_= 100 ms, τ_S_ = 50 ms) and used an interstimulus interval (ISI) of 100 ms. We extracted the sensory response in the maximally selective neuron to the stimulus in two simulations: 1) T2 present, 2) T2 absent. The simulation in Figure 6B was identical, except that the ISI was increased to 600 ms.

To quantify the effect of excitatory and suppressive time constants on the magnitude of response adaptation, we varied each of 𝜏_𝐸_ and 𝜏_𝑆_separately across 10 levels (0, 50, 100, 200, 300, 400, 500, 600, 700, and 800 ms) while keeping the other time constant fixed at zero, and assessed the sensory response across a range of ISIs (100 to 1500 ms, in 100 ms steps). To investigate whether temporal windows were required for response adaptation in this model, we also simulated a condition with 𝜏_𝐸_ and 𝜏_𝑆_both equal to zero. For each value of the time constants, we performed simulations where T1 was present or absent. To isolate the response to T2 when T1 was present, we followed previous work (Lisberger & Movshon, 1999; Priebe et al., 2002) by first subtracting out the activity elicited by a single stimulus (here, when T2 was absent), as follows:

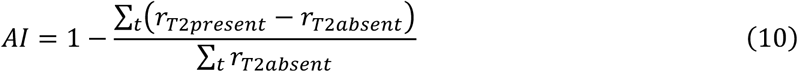

such that a higher adaptation index corresponds to a smaller isolated T2 response and thus stronger response adaptation. To examine how the time constants interact to affect response adaptation, we conducted additional simulations using combinations of shorter and longer time constants (𝜏_𝐸_ = 100 or 500 ms, 𝜏_𝑆_ = 50 or 500 ms), and computed suppression indices across the same range of ISIs (Figure 6E).

To assess the presence of response adaptation for non-identical stimuli (Figure 7), we performed further simulations in which we manipulated the difference in orientation between T1 (the adapting stimulus) and T2 (the test stimulus). In these simulations, we fixed the temporal window parameters (𝜏_𝐸_= 400 ms, 𝜏_𝑆_= 100 ms) and the ISI (100 ms) to focus on the effects of feature similarity and suppressive pooling on the magnitude of adaptation. To manipulate the tuning of the suppressive pool, we computed a pooling matrix that weighs the influence of one neuron on another when calculating suppressive drives, as follows:

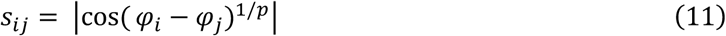

where 𝜑_𝑖_ and 𝜑_j_are the preferred orientations of the *i*th and *j*th neuron respectively, and *p* is a scaling parameter that determines the sharpness of the suppressive pooling. In the limit when *p* = ∞, all weights are 1, producing uniform suppressive pooling. In contrast, when *m* is small, suppression is mostly driven by similarly-tuned neurons, and when *p* = 0 neurons are only suppressed by themselves.

Across simulations, we fixed the test stimulus orientation at 0° and varied the adapting stimulus orientation in 10° steps from 0° (identical) to 90° (orthogonal). We separately adjusted the suppressive tuning across simulations at 7 levels (*p* = ∞, 1, 0.4, 0.2, 0.1, 0.04, 0). For each set of parameters, we measured the response in the neuron tuned to the test orientation in three separate simulations presenting: 1) the test stimulus alone (*r*_test_); 2) the adapting stimulus alone (*r*_adapt_); 3) both the adapting and test stimulus (*r*_both_). We then calculated the adaptation index, following Priebe and Lisberger (2002):

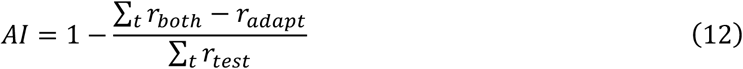

### Backward masking

Backward masking refers to the phenomena that perception of a stimulus can be impaired (“masked”) by a second stimulus presented shortly after the first (Breitmeyer & Öğmen, 2006; Enns & Di Lollo, 2000; Kovacs et al., 1995). We assessed backward masking in the model using a similar method as for response adaptation, except with simulations comparing the sensory response to T1 as a function of whether T2 was present vs absent:

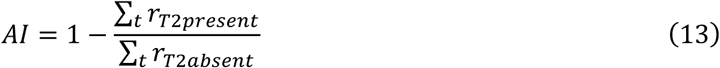

Again, we first conducted simulations using orthogonal stimulus orientations, with short time constants (τ_E_ = 100 ms, τ_S_ = 50 ms) and compared the effects of backward masking for stimulus onset asynchronies (SOA) of 250 ms (Figure 8A) and 500 ms (Figure 8B). We then quantified backward masking by calculating the response summed over time to T1 when T2 was present relative to when it was absent. We calculated this masking index across the full range of excitatory and suppressive time constants and SOAs (Figures 7C-D). Finally, we conducted additional simulations assessing the interaction of the two time constants on backward masking using combinations of one shorter and one longer time constant (τ_E_ = 100 or 500 ms, τ_S_= 50 or 500 ms), and computed masking indices across the same range of SOAs (Figure 8E).

### Contrast-dependent stimulus interactions

We assessed how the contrast of one stimulus affects the response to the other stimulus through temporal normalization. For initial simulations, we used fixed time constants (𝜏_𝐸_ = 400 ms, 𝜏_𝑆_= 100 ms), with the SOA (250 ms) and stimulus contrasts (64% vs. 16%) chosen to match the previous behavioral findings. We performed simulations independently manipulating the contrast of both T1 and T2 (64% or 16% contrast level), and extracted model performance for each target. We first examined how the model responses to each target at high contrast was affected by the contrast of the non-target (Figures 8A-B). We then computed the model’s behavioral discrimination performance (measured as d’) from the output of the decision layer, based on recent empirical findings showing that a high- vs. low-contrast non-target stimulus presented before or after a target stimulus can result in reduced perceptual discriminability of the target (Epstein & Denison, 2023). Model responses were scaled to produce d’ values closer to behavioral estimates (s_T1_ = s_T2_ = 1×10^4^). Model d’ was calculated for each target stimulus based on the target and non-target contrast (Figure 9C).

To assess how contrast-dependent suppression for T1 and T2 was affected by the excitatory and suppressive temporal windows, we performed further simulations in which we varied 𝜏_𝐸_ and 𝜏_𝑆_from 0 to 1000 ms in 50 ms steps. For each simulation, we measured the model d’ for each target stimulus (T1 and T2) as a function of the contrast of the non-target (NT), as above. We then calculated a contrast-dependent suppression index for each stimulus as:

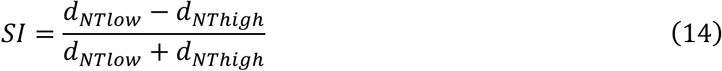

To identify values of 𝜏_𝐸_ and 𝜏_𝑆_that produced suppression in both T1 and T2, we calculated a joint suppression index across the two targets by multiplying the suppression indices of T1 and T2 for each parameter combination. This joint index is maximized when a specific combination of 𝜏_𝐸_ and 𝜏_𝑆_results in contrast-dependent suppression in both stimuli, but not when one or both stimuli show little- to-no suppression.

## Acknowledgements

This work was supported by Boston University startup funds to R.N.D. We thank the members of the Denison Lab for their feedback on the project, particularly Michael Epstein and Karen Tian for helpful discussions.

